# Quantitative nascent proteome profiling by dual pulse labeling with *O*-propargyl-puromycin and stable isotope labeled amino acids

**DOI:** 10.1101/2020.06.06.137521

**Authors:** Junki Uchiyama, Yasushi Ishihama, Koshi Imami

## Abstract

Monitoring translational regulation in response to environmental signals is crucial for understanding cellular proteostasis. However, only limited approaches are currently available for quantifying acute changes in protein synthesis induced by stimuli. Recently, a clickable puromycin analog, *O*-propargyl-puromycin (OPP), was developed and applied to label the C-termini of nascent polypeptide chains (NPCs). Following affinity purification via a click reaction, OPP allows for a proteomic analysis of NPCs. Despite its advantage, the affinity purification of NPCs using magnetic beads or resins inherently suffers from significant non-specific protein binding, which hinders accurate quantification of the nascent proteins. To address this issue, we employed dual pulse labeling of NPCs with both OPP and stable isotope labeled amino acids to distinguish *bona fide* NPCs from non-specific proteins, thereby enabling the accurate quantitative profiling of NPCs. We applied this method to dissecting the transcription-coupled translation responses and quantified ~3,000 nascent proteins. We found that the translation of a subset of ribosomal proteins (e.g., RPSA, RPLP0) as well as signaling proteins (e.g., BCAR3, EFNA1, DUSP1) was significantly repressed by transcription inhibition. Together, the present method provides an accurate and broadly applicable nascent proteome profiling for many biological applications at the level of translation.

## Introduction

Protein synthesis is an essential biological process in all living organisms to maintain cellular proteostasis. Translational regulation through initiation factors (e.g., 4E-BP, eIF2), protein kinases (e.g., mTOR, p70 S6 kinase), ribosomes as well as the *cis*-regulatory elements of mRNAs can dynamically change a proteome profile in order to adapt to environmental cues rapidly (1). Therefore, monitoring a nascent proteome profile is key to understanding gene expression control and subsequent cellular responses.

Mass spectrometry (MS)-based proteomics can be used to monitor proteome dynamics. Pulse stable isotope labeling by amino acids in cell culture (pSILAC) (2, 3) and quantitative noncanonical amino acid tagging (QuaNCAT) (4, 5) utilize stable isotope labeled amino acids and/or a bioorthogonal noncanonical amino acid (6) that are incorporated into nascent proteins during mRNA translation. Hence, the labeled newly synthesized proteins can be distinguished from non-labeled pre-existing proteins, thereby allowing a global quantification of the translation products with MS. These methods allow for quantifying ‘young’ proteins that are newly synthesized during the pulse labeling, but they cannot exclusively capture nascent polypeptide chains (NPCs) being elongated by the ribosomes. To profile NPCs, puromycin or its analogs have been applied to labeling NPCs, as they are tRNA analogs and can be incorporated into the C-termini of NPCs during the elongation of polypeptide chains (7). Using a puromycin analog, Aviner *et al*. developed puromycin-associated nascent chain proteomics (PUNCH-P) (8). PUNCH-P relies on *in vitro* cell-free labeling of NPCs with biotin-puromycin, which then enables the isolation of biotin-labeled nascent proteins with streptavidin beads. As an alternative approach, Barrett *et al*. and Forester *et al*. established methods for nascent proteome analysis using *O*-propargyl puromycin (OPP) (9, 10). A cell-permeable OPP can label nascent elongating polypeptides *in vivo*, which are subsequently conjugated to biotin azide molecules using click chemistry, and the biotinylated NPCs are captured with streptavidin beads. While this method was shown to be a powerful tool for profiling nascent polypeptide *in vivo*, using streptavidin beads inherently suffers from non-specific protein binding as in any affinity purifications (11), hindering the detection of *bona fide* nascent proteins.

In this study, we devised a method for accurate quantitative nascent proteome profiling by combining the pulse labeling of OPP and SILAC amino acids. We demonstrate that our approach successfully separates nascent proteins and non-specific binders, allowing more accurate quantitative profiling than existing methods. Using this method, we assessed translational responses coupled with transcription and uncovered proteins modulated at the level of protein synthesis.

## Materials and Methods

### Cell culture and pulse labeling

HeLa cells obtained from ATCC were cultured in Dulbecco’s Modified Eagle Medium (DMEM) media (Fujifilm Wako, Osaka, Japan) supplemented with 10% FBS (Thermo Fisher Scientific, Waltham, USA) and 100 μg/mL kanamycin. For pulse labeling, the cell culture medium was switched to arginine- and lysine-free DMEM (Thermo Fisher Scientific, Waltham, USA) supplemented with 30 μM *O*-propargyl-puromycin (OPP) (Jena Bioscience, Jena, Germany), 10% FBS and either “heavy” amino acids [0.398 mM L-(^13^C_6_,^15^N_4_)-arginine (Arg“10”) and 0.798 mM L-(^13^C_6_,^15^N_2_)-lysine (Lys“8”)] or “medium” amino acids [0.398 mM L-(^15^N_4_)-arginine (Arg“4”) and 0.798 mM L-(D_4_)-lysine (Lys“4”)] (Cambridge Isotope Laboratories, Tewksbury, USA), and incubated for 2 h. For actinomycin D (actD) treatment, HeLa cells were first pre-incubated with 50 nM actD for 2 h, and subsequent pulse labeling with OPP and SILAC amino acids was performed in the presence of 50 nM actD as described above. Of note, the effect of “light” amino acids (e.g., derived from FBS or by recycling) on SILAC labeling was negligible (Supplementary Fig. 1A), consistent with our previous report (12). All cells were maintained in a humidified 37°C incubator with 5% CO_2_.

### Click reaction of nascent proteins with biotin-puromycin

We followed a protocol for the click reaction described elsewhere (9) with a slight modification. Briefly, HeLa cells were lysed with a buffer [100 mM HEPES-NaOH (pH 7.5), 150 mM NaCl, 1% Nonidet P-40 (NP-40), 1% protease inhibitor], and cell debris was removed by centrifugation (4°C, 16,000 x g, 30 min). The protein concentration was measured using a BCA assay (Thermo Fisher Scientific), and 700 μg/600 μL protein per sample was used for the proteomic analysis. For the click reaction, a reagent mix was first prepared by adding stock solutions to achieve 6.25% SDS, 0.625 mM biotin-azide (Thermo Fisher Scientific), 6.25 mM TCEP (tris(2-carboxyethyl)phosphine) (Merck Millipore, Burlington, USA), 0.625 mM TBTA (Tris[(1-benzyl-1H-1,2,3-triazol-4-yl)methyl]amine) (Merck Millipore), and 6.25 mM CuSO_4_ (Merck Millipore). The reagent mix was then added to the cell lysates (1:5.25, vol/vol) to achieve 1% SDS, 100 μM biotin-azide, 1 mM TCEP, 100 μM TBTA, and 1 mM CuSO_4_, and the sample was incubated for 1.5 h at room temperature.

### Affinity purification of biotinylated proteins using streptavidin beads

After the click reaction, proteins were precipitated by adding 5 volumes of cold acetone and stored overnight (−20°C). The precipitated proteins were pelleted by centrifugation at 3,500 × *g* and 4°C for 5 min and washed twice with 500 μL cold methanol. Protein pellets were resuspended in 120 μL PBS containing 1% SDS and desalted by passing through Amicon Ultra 0.5 mL 10K cutoff desalting columns (Merck Millipore) equilibrated with 1% NP-40, 0.1% SDS in PBS. Biotinylated proteins were affinity purified with magnetic streptavidin beads (Thermo Fisher Scientific) (40 μL slurry per sample) at 4°C overnight with slow rotation. Beads were washed twice with 0.2 mL 1% NP-40, 0.1% SDS in PBS for 10 min, three times with 0.2 mL ice-cold 6 M urea in PBS for 15 min, and three times with 0.2 mL ice-cold PBS for 15 min, all at 4 °C with slow rotation. Finally, the beads were rinsed with 50 mM ammonia bicarbonate.

### On beads digestion, desalting and fractionation of peptides

The samples were incubated with 10 mM DTT at 37°C for 30 min to reduce the disulfide bonds of the proteins, followed by alkylation with 50 mM iodoacetamide at 37°C for 30 min in the dark. The proteins were digested with 500 ng trypsin (Promega, Madison, USA) overnight at 37°C on a shaking incubator. Tryptic digestion was quenched by adding 0.5% TFA (final concentration). The resulting peptides were desalted and fractionated with a SDB-XC-SCX StageTip (13). The sample solution was evaporated by SpeedVac and then resuspended in 0.5% TFA and 4% ACN for LC/MS/MS.

### LC/MS/MS analysis

Nano-scale reversed-phase liquid chromatography coupled with tandem mass spectrometry (nanoLC/MS/MS) was performed by an Orbitrap Fusion Lumos mass spectrometer (Thermo Fisher Scientific), connected to a Thermo Ultimate 3000 RSLCnano pump and an HTC-PAL autosampler (CTC Analytics, Zwingen, Switzerland) equipped with a self-pulled analytical column (150 mm length × 100 μm i.d.) (14) packed with ReproSil-Pur C18-AQ materials (3 μm, Dr. Maisch GMBH, Ammerbuch, Germany). The mobile phases consisted of (A) 0.5% acetic acid and (B) 0.5% acetic acid and 80% acetonitrile. Peptides were eluted from the analytical column at a flow rate of 500 nL/min by altering the gradient: 5-10% B in 5 min, 10-40% B in 60 min, 40-99% B in 5 min and 99% for 5 min. The Orbitrap Fusion Lumos instrument was operated in the data-dependent mode with a full scan in the Orbitrap followed by MS/MS scans for 3 sec using higher-energy collisional dissociation (HCD). The applied voltage for ionization was 2.4 kV. The full scans were performed with a resolution of 120,000, a target value of 4×10^5^ ions and a maximum injection time of 50 ms. The MS scan range was *m/z* 300–1,500. The MS/MS scans were performed with a 15,000 resolution, a 5×10^4^ target value and a 50 ms maximum injection time. The isolation window was set to 1.6, and the normalized HCD collision energy was 30. Dynamic exclusion was applied for 20 sec.

### Database searching and protein quantification

All raw files were analyzed and processed by MaxQuant (v1.6.2.10) (15). Search parameters included two missed cleavage sites and variable modifications such as L-(^13^C_6_,^15^N_4_)-arginine, L-(^13^C_6_,^15^N_2_)-lysine, L-(^15^N_4_)-arginine, L-(D_4_)-lysine, methionine oxidation, and protein N-terminal acetylation. Cysteine carbamidomethylation was set as a fixed modification. The peptide mass tolerance was 4.5 ppm, and the MS/MS tolerance was 20 ppm. The database search was performed with Andromeda (16) against human SwissProt Database (version 2017-04, 20,199 protein entries) with common contaminants and enzyme sequences. The false discovery rate (FDR) was set to 1% at the peptide spectrum match (PSM) level and protein level. For the SILAC-based protein quantification related to Fig. 2, a minimum of one ratio count (unique peptide ion) was set, and the ‘re-quantify’ and ‘match between runs’ functions were employed. Raw H/M and H/L ratios were used for quantification. For the quantification of nascent proteins induced by actD (related to Fig. 3), a minimum of three ratio counts for H/M ratios was used and the other parameters were the same as above. Normalized H/M ratios were used to correct a mixing error between the “H” and “M” lysates. InstantClue (17) was used to generate multi scatter plots and boxplots.

### Repository for LC/MS/MS raw data

The proteomics data have been deposited to the ProteomeXchange Consortium via the jPOST (18, 19) partner repository with the dataset identifier JPST000860 (PXD019459 for ProteomeXchange).

### Computing protein abundance using iBAQ

The intensity-based absolute quantification (iBAQ) algorithm (20) computes the sum of all peptides intensities divided by the number of theoretically observable peptides, which provides a rough estimation of protein abundance within a sample. iBAQ intensities from the “heavy” channel were used as a relative protein abundance of NPCs within the sample and compared to the read counts of ribosome protected fragments (RPFs) obtained from the ribosome profiling (21).

### Comparisons with ribosome profiling data

Ribosome profiling data were obtained from a previously published data set (21) in which cell cycle-dependent (that is, G1, S, M) translational changes were analyzed in HeLa cells. Transcripts having read counts within 0-10 were eliminated. For our proteomic results, we only used proteins showing a magnitude change of at least 2-fold (i.e., H/M ratio>2) as those proteins are more likely to be *bona fide* nascent proteins. To compare our proteomic result (asynchronous HeLa cells) with the ribosome profiling data, we used data from the G1 cell cycle stage because G1 is the major cell cycle phase in asynchronous cells. Another reason is that overall protein synthesis is not affected by the cell cycle phase (3, 22).

### Western blotting

HeLa cells were lysed with lysis buffer (100 mM HEPES-NaOH pH 7.5, 1% NP-40, 150 mM NaCl and 1% protease inhibitor). Supernatants after centrifugation (16,000 g, 30 min, 4°C) were biotinylated by a click reaction as described above. The samples were re-suspended in LiDS loading sample buffer (Invitrogen) with 50 mM DTT and incubated at 70°C for 5 min. The protein samples were loaded onto a 4–12% gradient SDS-polyacrylamide gel (Thermo Fisher Scientific) and separated using electrophoresis. The proteins were then transferred to a PVDF membrane (Merck Millipore) using a semi-dry western blot transfer system set to a constant current of 200 mA for 30 min. The membranes were first blocked by incubating in 5% (w/v) BSA in Tris-buffered saline and 0.1% tween (TBS-tween) and then incubated with HRP-conjugated Streptavidin (Thermo Fisher Scientific) diluted 1:50,000 in 5% BSA in TBS-tween for 4 h while rotating at room temperature. Membranes were washed three times in TBS-tween and developed with ECL reagent (Thermo Fisher Scientific).

### Click reactions for fluorescence nascent RNA imaging

HeLa cells were incubated with 1 mM 5-ethylene uridine (baseclick, Neuried, Germany) in the presence of DMSO or actinomycin D (Fujifilm Wako) for 4 h. The cells were washed with PBS and fixed with 1% paraformaldehyde in PBS for 10 min at room temperature. Click reactions were performed as described elsewhere (3, 23). Briefly, the cells were permeabilized with 1% BSA with 0.2% saponin in PBS for 15 min and then incubated with click solution (1 μM Chromeo™ 488-Azide (baseclick), 10 mM (+)-Sodium L-ascorbate and 2 mM CuSO_4_) for 30 min at room temperature. The cells were washed once with 1% BSA in PBS before imaging.

## Results

### Experimental design

OPP is a clickable tRNA analog that can be incorporated into the C-termini of elongating NPCs (9). After treating cells with OPP, OPP-labeled NPCs can be biotinylated via a click reaction, followed by affinity purification of the biotinylated NPCs using streptavidin beads (Fig. 1A). One critical technical issue in such affinity purification is that a dozen proteins can be bound to the beads in a non-specific manner (11), thereby hindering the identification and quantification of the nascent proteome. It is, therefore, crucial to separate NPCs and non-specific binding proteins. We reasoned that dual pulse labeling with both OPP and SILAC amino acids should allow us to solve this issue; SILAC amino acids should be in principle incorporated into NPCs during translation in addition to OPP, which results in discriminating “heavy” labeled-NPCs and non-specific “light” (non-labeled) proteins (Fig. 1A).

**Figure 1.**
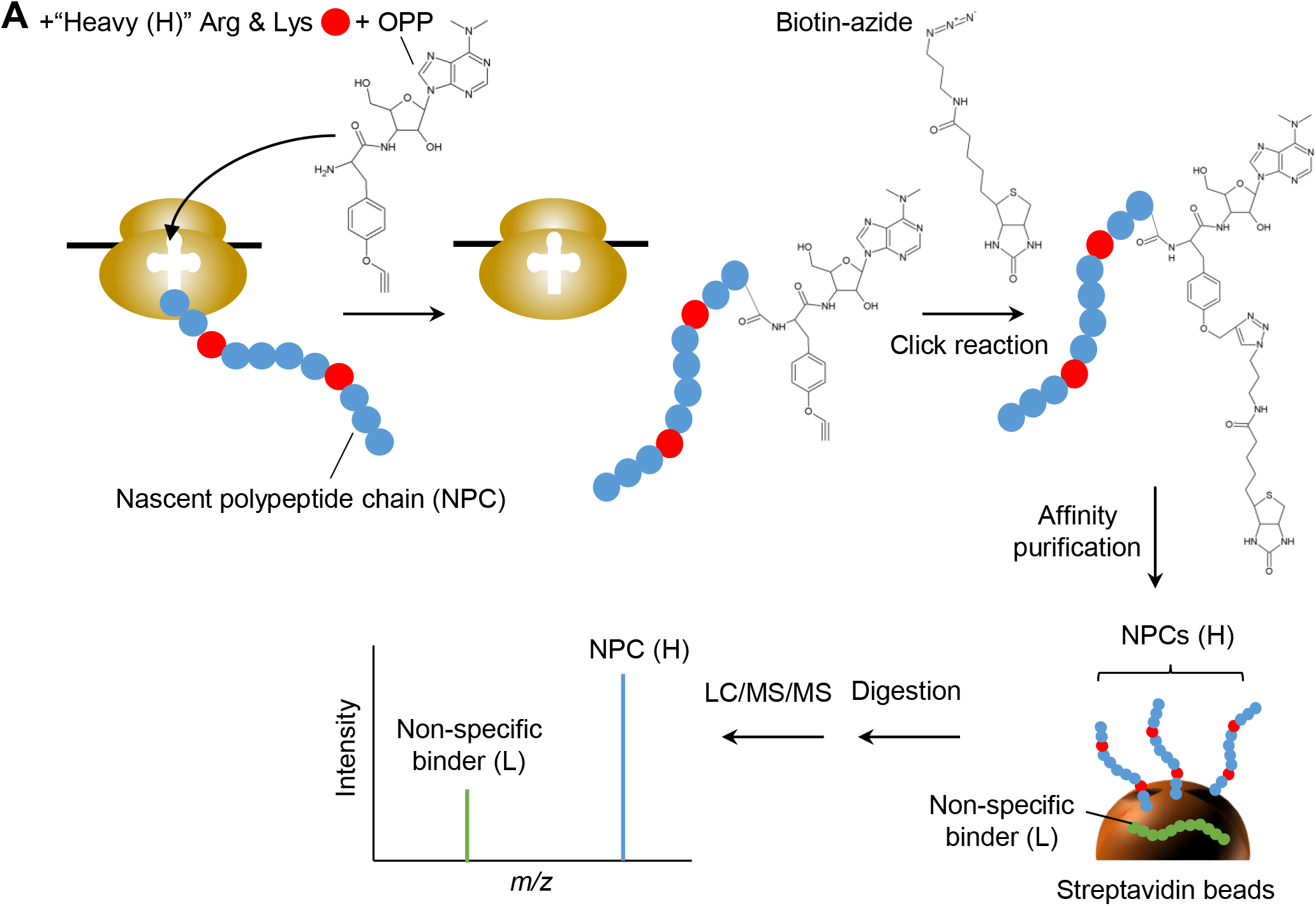
Experimental design. (A) Concept of the method. Cells are pulse-labeled with OPP and SILAC amino acids (“heavy” arginine and lysine). Proteins are then extracted, biotinylated via click reaction, affinity captured with streptavidin beads, and digested into peptides. The resulting peptide sample is analyzed by LC/MS/MS. Since nascent proteins are labeled with “heavy” amino acids, nascent proteins (“heavy”) and non-specific proteins (“light”) can be separated with MS.

### Nascent proteome profiling by combining pulse OPP and SILAC labeling

First, we confirmed using western blotting that our experimental setup detected a reasonable amount of OPP- and biotin-labeled proteins in HeLa cells while neither omitting OPP nor the inhibition of protein synthesis by cycloheximide (CHX) treatment (Fig. 2A). These results indicate that nascent proteins can be labeled with OPP in a translation-dependent manner.

**Figure 2.**
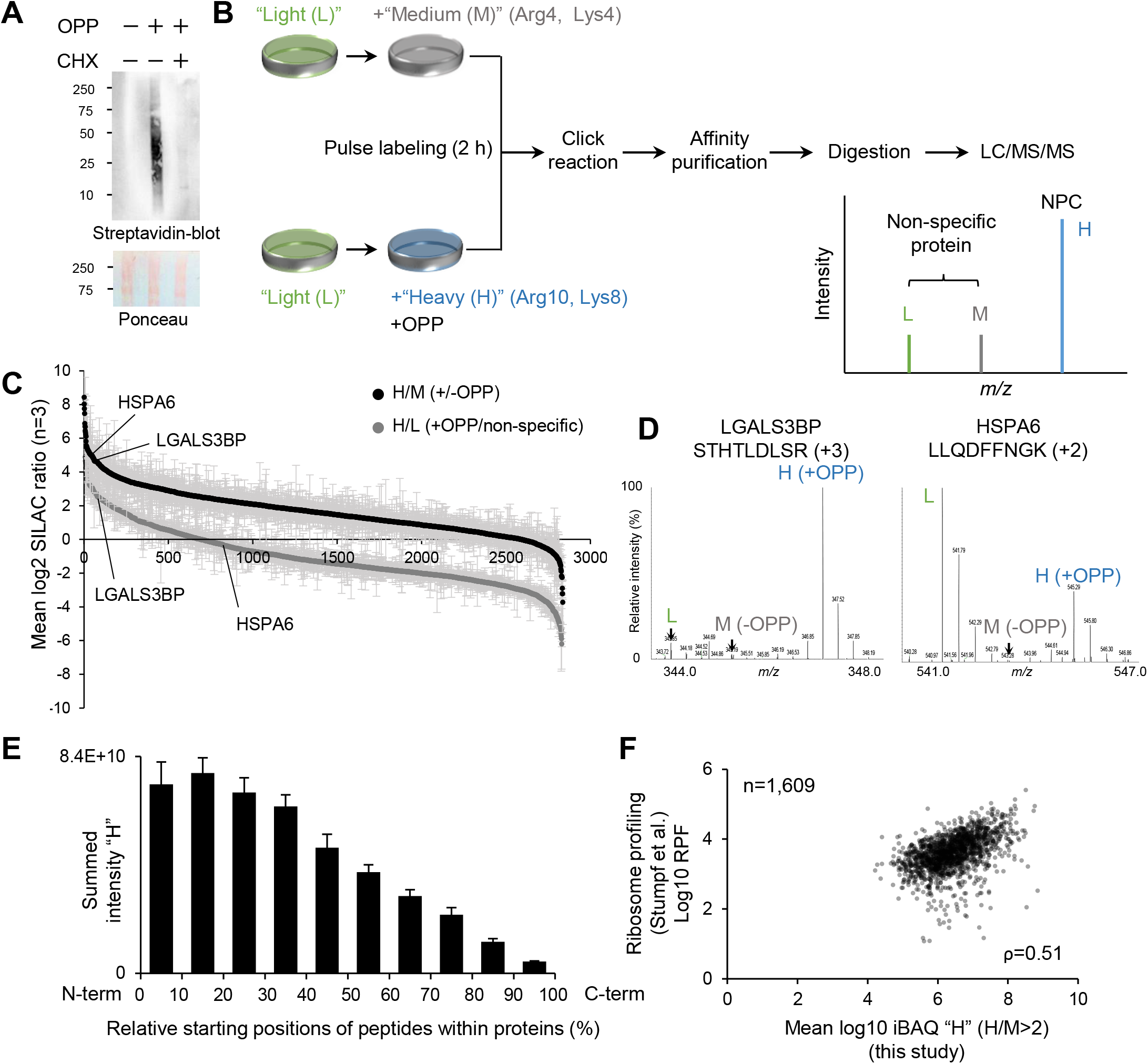
Demonstration of nascent proteome profiling. (A) (Top) Western blots and (bottom) Ponceau staining of whole cell lysates from HeLa cells treated with OPP, OPP and CHX, or neither. OPP-labeled proteins were biotinylated through click chemistry, and biotinylated proteins were detected with HRP-conjugated streptavidin. (B) Experimental scheme for the proteomic analysis. HeLa cells were pulse labeled with OPP and “heavy” SILAC amino acids (H: Arg10, Lys8) or only “medium” SILAC amino acids (M: Arg4, Lys4). The subsequent procedures are the same as in Fig. 1. Proteins labeled with “heavy” amino acids (i.e., high H/M ratios) represent nascent proteins. (C) An overview of OPP-treated(“H”)/control(“M”) and OPP-treated(“H”)/non-specific(“L”) ratios (log2 scale), plotted from largest to smallest. The means (±S.D.) of ratio were determined from at least three independent experiments. (D) Exemplary MS spectra for two peptides; one for STHTLDLSR (LGALS3BP) and one for LLQDFFNGK (HSPA6). The isotope clusters of the L, M, and H peaks correspond to non-specific binding proteins from the pre-existing proteome pool (“L”), and from the “medium”-labeled nascent proteome (“M”) and *bona fide* NPCs (“H”), respectively. (E) Relative starting positions of identified peptides within proteins. Only proteins that exhibit H/M>2 were used for this analysis. (F) iBAQ intensities from individual proteins modestly correlated with corresponding RPF read counts from individual genes. The ribosome profiling data were obtained from a previous study (21).

To demonstrate a proof-of-concept for proteomic application, we first pulse-labeled HeLa cells with both OPP and “heavy” amino acids (H: Arg”10”, Lys”8”) for 2 h (Fig. 2B). As a negative control, the cells were treated with only “medium” amino acids (M: Arg”4”, Lys”4”) and OPP was omitted. Following the established protocol (see Material and Methods) which includes protein extraction, click reaction, isolation of NPCs, and digestion to peptides, the peptide mixtures were subjected to nanoLC/MS/MS. In total, 2,837 proteins were quantified in all three independent experiments (Fig. 2C and Supplementary Table 1). The MS spectra of representative examples (STHTLDLSR and LLQDFFNGK from LGALS3BP and HSPA6, respectively) are shown in Fig 2D. We observed the isotope clusters of the light, medium and heavy peaks that correspond to non-specific binding proteins from the pre-existing proteome pool (“light”: L) and non-specific binding proteins from the “medium”-labeled nascent proteome (M) and *bona fide* NPCs (H), respectively. Hence, the observed SILAC ratios (H/M and H/L) can be used to assess how selectively NPCs could be enriched with this method. As shown in Fig. 2D, we observed a higher enrichment of “heavy” NPCs in comparison to “medium” proteins. Indeed, we overall observed a trend of high H/M ratios, which corresponds to 93% (H/M>1) and 69% (H/M>2) of total quantified proteins (Fig. 2C). This result indicates that OPP-labeled NPCs could be captured and quantified successfully with this approach. Of note, we observed lower H/L ratios compared with H/M ratios (Fig. 2C and 2D right), which indicates a substantial amount of non-specific proteins bound to the beads. Reproducibility of SILAC ratios between replicates was overall high (Pearson correlation coefficient r>0.72 and r>0.91 for H/M and H/L ratios, respectively) (see also Supplementary Fig. 1B). However, the weaker correlation was observed for H/M ratios possibly due to lower signal intensities of “medium”- and “heavy”-proteins compared with “light” proteins (Supplementary Fig. 1C).

These results demonstrated that our method enables the separation of NPCs and non-specific binding proteins, thus providing accurate nascent proteome profiling.

### OPP- and heavy-labeled proteins show expected features of nascent polypeptide chains

To further validate whether the “heavy”-labeled proteins showing at least H/M>2 exhibits a feature of NPCs, we performed an additional data analysis. Ribosomes elongate polypeptides from their amino (N-) terminal end to their carboxyl (C-) terminal end, and OPP causes premature termination of the elongation of polypeptides. Hence, positions of “heavy”-labeled peptides identified by this method are expected to show a bias towards the N-termini of corresponding proteins. As expected, the identified peptides were biased towards the N-termini of proteins (Fig. 2E), while the less pronounced trends were observed for the proteins showing H/M<2 (Supplementary Fig. 1D left, center). Besides, no such trend was observed for proteins identified from a whole cell lysate (Supplementary Fig. 1D right). This analysis supports our conclusion that “heavy” proteins (at least H/M>2) are the most likely to be NPCs.

We next compared our result with a ribosome profiling data. Ribosome profiling is a next-generation sequencing (NGS)-based method (24) that sequences mRNA fragments bound to ribosomes (RPFs), as an indicator of the levels of translation of each mRNA region. The present MS-based method and ribosome profiling quantify distinct molecules (*i.e*., proteins and nucleic acids) and give distinct readouts (*i.e*., MS signal intensity and NGS read counts); however, given that both methods provide a snapshot of how actively individual mRNAs are translated, the MS signal intensities of NPCs should correlate with read counts of RPFs. To test this, we used intensity-based absolute quantification (iBAQ) (20), which computes the relative protein abundance within a sample based on MS signal intensities normalized by the number of theoretically observable peptides per protein. We then compared the averaged iBAQ values of individual “heavy” NPCs showing at least H/M>2 with read counts of RPFs of individual genes obtained from previously published data (21). As expected, we observed a positive correlation (Spearman’s rho = 0.51) between our proteomic result and the ribosome profiling data (Fig. 2F). We also found that the correlation of iBAQ values between replicates was high (r>0.94) (Supplementary Fig. 1E). Note that iBAQ “L” and “M” also exhibited a positive correlation (Supplementary Fig. 1F left, center) as the degree to which proteins bind to beads non-specifically depends on their cellular protein abundance (Supplementary Fig. 1F right). On the one hand, we used only high confident NPCs that showed H/M>2 (see also Fig. 2E) to check the correlation with the ribosome profiling, indicating that “heavy” intensities are the most likely to reflect protein abundance from an NPC pool. Together, this finding suggests that our method recapitulates the ribosome profiling data and that the overall protein synthesis rate is likely to be consistent between the two methods.

### Monitoring translational responses coupled with transcription

Having established an experimental scheme for nascent proteome profiling, we next sought to apply this method to assessing the transcription-translation relationship. It is known that transcription by RNA polymerase (RNAP) is coupled to translation by the ribosome in bacteria via formation of an “expressome” complex of the RNAP and ribosome (25). However, little is known about the transcription-translation relationship in human cells in which ribosomes and RNAPs are spatially compartmentalized. We thus sought to monitor translational responses upon transcriptional inhibition. To this end, we treated HeLa cells with actinomycin D (actD) which inhibits RNAP I potently as well as RNAP II and III (26). Indeed, 50 nM actD was sufficient to inhibit RNAP activity, as judged by the attenuated fluorescent signals of nascent RNA compared with the control condition (Fig. 3A)

**Figure 3.**
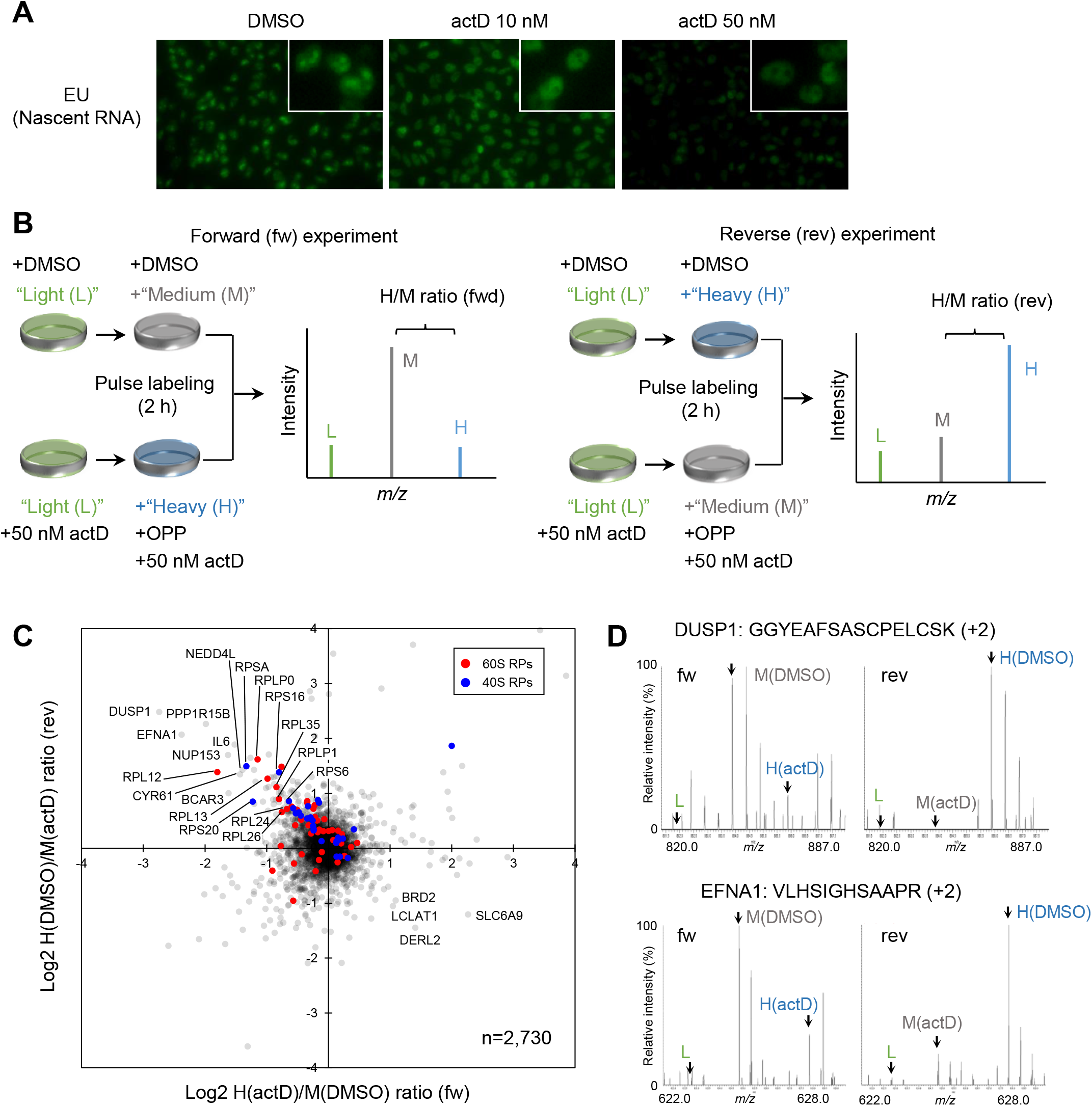
Profiling the nascent proteome in response to actinomycin D. (A) Visualizing and quantifying nascent RNA by EU (5-ethynyl uridine) labeling and click chemistry detection. Nascent RNAs were labeled with a clickable EU in the presence of actD (0, 10, 50 nM) and visualized through a click reaction with a green fluorescent dye. (B) Experimental scheme. HeLa cells were treated with 50 nM actD or DMSO for 2 h and then pulse labeled with OPP and SILAC amino acids for another 2 h. Two independent experiments were performed based on “label-swap” replicates in which forward (fw) and reverse (rev) experiments represent “H” actD + “M” DMSO and “M” actD + “H” DMSO, respectively. H/M ratios indicate the difference in protein synthesis between the two conditions (actD v.s. DMSO). (C) A scatter plot showing log2 H/M ratios obtained by the forward and reverse experiments. Ribosomal proteins are highlighted by red (60S ribosomal proteins) and blue (40S ribosomal proteins). (D) Exemplary MS spectra for two peptides; one for GGYEAFSASCPELCSK (DUSP1) and one for VLHSIGHSAAPR (EFNA1).

For the proteome analysis, we first pre-incubated HeLa cells in the presence of either 50 nM actD or vehicle (DMSO) for 2 h and then added OPP and SILAC amino acids for an additional 2 h (Fig. 3B). We pooled differentially labeled actD and DMSO lysates together for further proteomic analysis (“H” actD + “M” DMSO in forward (fw) and “M” actD + “H” DMSO in reverse (rev) experiments). The fw and rev experiments represent “label swap” replicates that eliminate biases introduced by the labeling procedure. The H/M (fw) and M/H (rev) ratios for each protein represent the relative amounts of NPCs in actD compared with DMSO.

Using this approach, we quantified 3,967 proteins in both duplicates, of which 2,730 proteins were quantified from at least 3 unique peptide ions and used for further analysis (Fig. 3C and Supplementary Table 2). Exemplary MS spectra for two peptides from DSUP1 and EFNA1, are shown in Fig. 3D. We observed that the protein synthesis of a subset of ribosomal proteins (RPs), including RPSA, RPLP0, and RPL12, was significantly inhibited by the actD treatment. This is in line with actD blocking rRNA synthesis through the inhibition of RNAP I (27), and therefore the translation of those RPs might be inhibited as the RPs lost their assembly partners (*i.e*., rRNA-RP subcomplexes). Interestingly, only a specific subset of RPs was translationally down-regulated (see Discussion). Moreover, we observed that several signaling proteins (BCAR3, CYR61, IL6, EFNA1, DUSP1, and PPP1R15B) showed translational repression of a magnitude change of at least 1.5-fold in both replicates. In contrast, the protein synthesis of other proteins, such as DERL2 and LCLAT1, was elevated. Because actD affects transcription, we checked the genomic regions for the regulated proteins but found no region specific-regulation (Supplementary Fig. 2A). Collectively, we successfully quantified acute changes in protein synthesis induced by actD, and uncovered transcription-coupled translation products including RPs and signaling proteins.

### Comparison with a conventional pulse SILAC approach

We next sought to compare the present method with a conventional pSILAC approach. While our method allows for quantifying acute changes (within 2 h in this study) in protein synthesis, pSILAC requires longer pulse labeling time (typically >12 h) to detect “heavy”-labeled proteins. Therefore, we can not directly compare the two methods under the same labeling condition. To this end, we performed pSILAC experiments in which HeLa cells were pulse-labeled with “heavy” or “medium” amino acids for 4 h to obtain a detectable level of “heavy” and “medium” proteins and to minimize the labeling time as much as possible (Fig. 4A).

**Figure 4.**
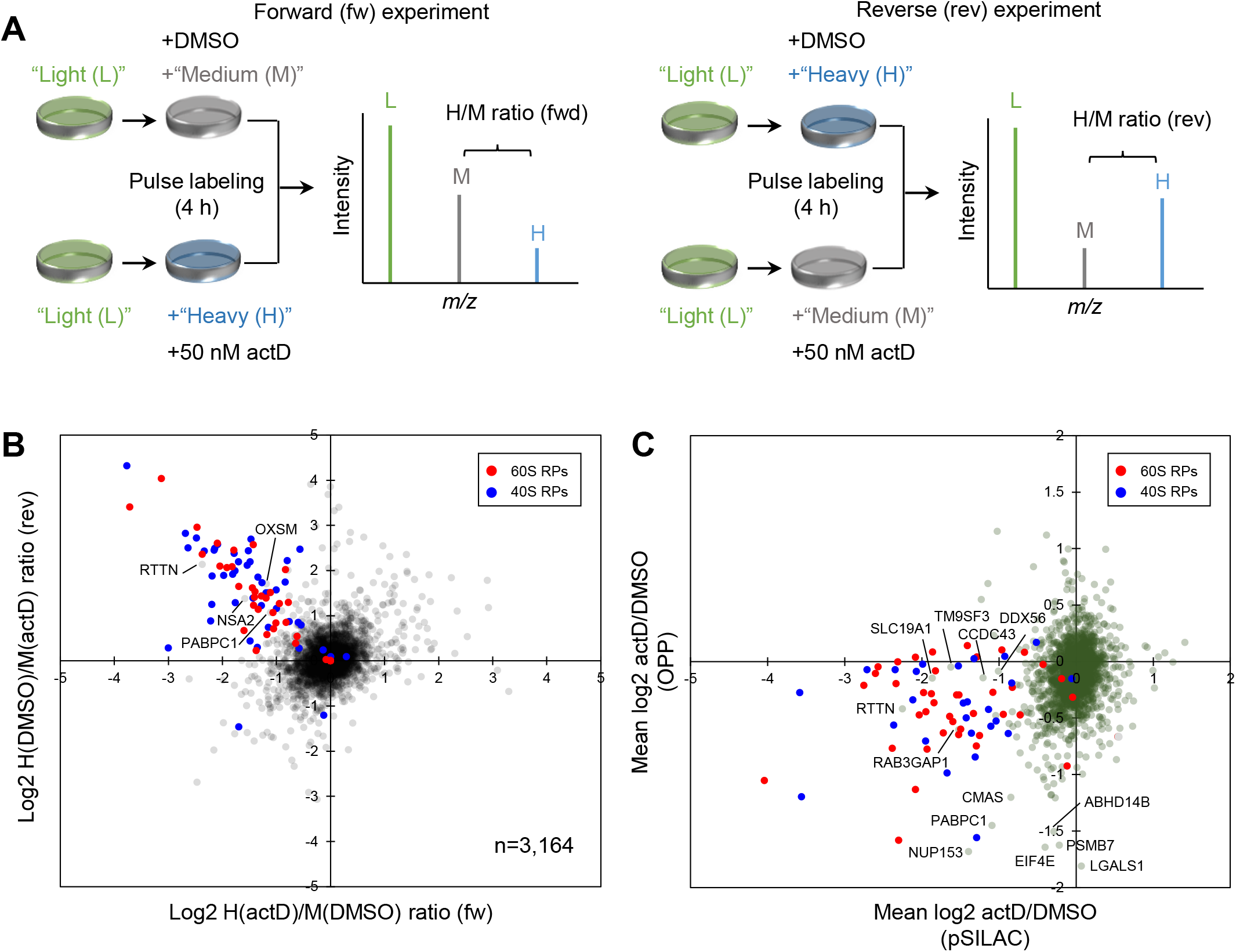
pSILAC approach for profiling the newly synthesized proteome in response to actinomycin D. (A) Experimental scheme. HeLa cells were pulse labeled with OPP and SILAC amino acids for 4 h in the presence of either 50 nM actD or DMSO. Two independent experiments were performed based on “label-swap” replicates in which forward (fw) and reverse (rev) experiments represent “H” actD + “M” DMSO and “M” actD + “H” DMSO, respectively. H/M ratios indicate the difference in protein synthesis between the two conditions (actD v.s. DMSO). (B) A scatter plot showing log2 H/M ratios obtained by the forward and reverse experiments. Ribosomal proteins are highlighted by red (60S ribosomal proteins) and blue (40S ribosomal proteins). (C) A scatter plot showing log2 actD/DMSO ratios obtained by the present nascent proteome profiling (Fig. 3C) and the pSILAC (Fig. 4B).

In total, 3,164 proteins were quantified in both duplicates and from at least 3 unique peptide ions (Fig. 4B and Supplementary Table 2). Consistent with our results (Fig. 3C), RPs were among the most impacted proteins in the pSILAC (Fig. 4B). Perhaps not surprisingly, we observed the differences in quantitative changes between the two methods (Fig. 4C). For example, while our method captured rapid changes in the translation of only a specific subset of the RPs within 2 h, most of the quantified RPs appear to be already modulated within 4 h in the pSILAC experiment. Notably, secreted proteins and specific regulatory proteins, including transcriptional factors, E3 ubiquitin ligases, and signaling proteins, were exclusively identified and quantified in our approach (Table 1). One of the advantages in our method is that even secreted proteins can be captured by enriching their nascent forms in the cytoplasm, which is in marked contrast to the conventional pSILAC approach that mainly captures intracellular proteomes.

**Table 1:**
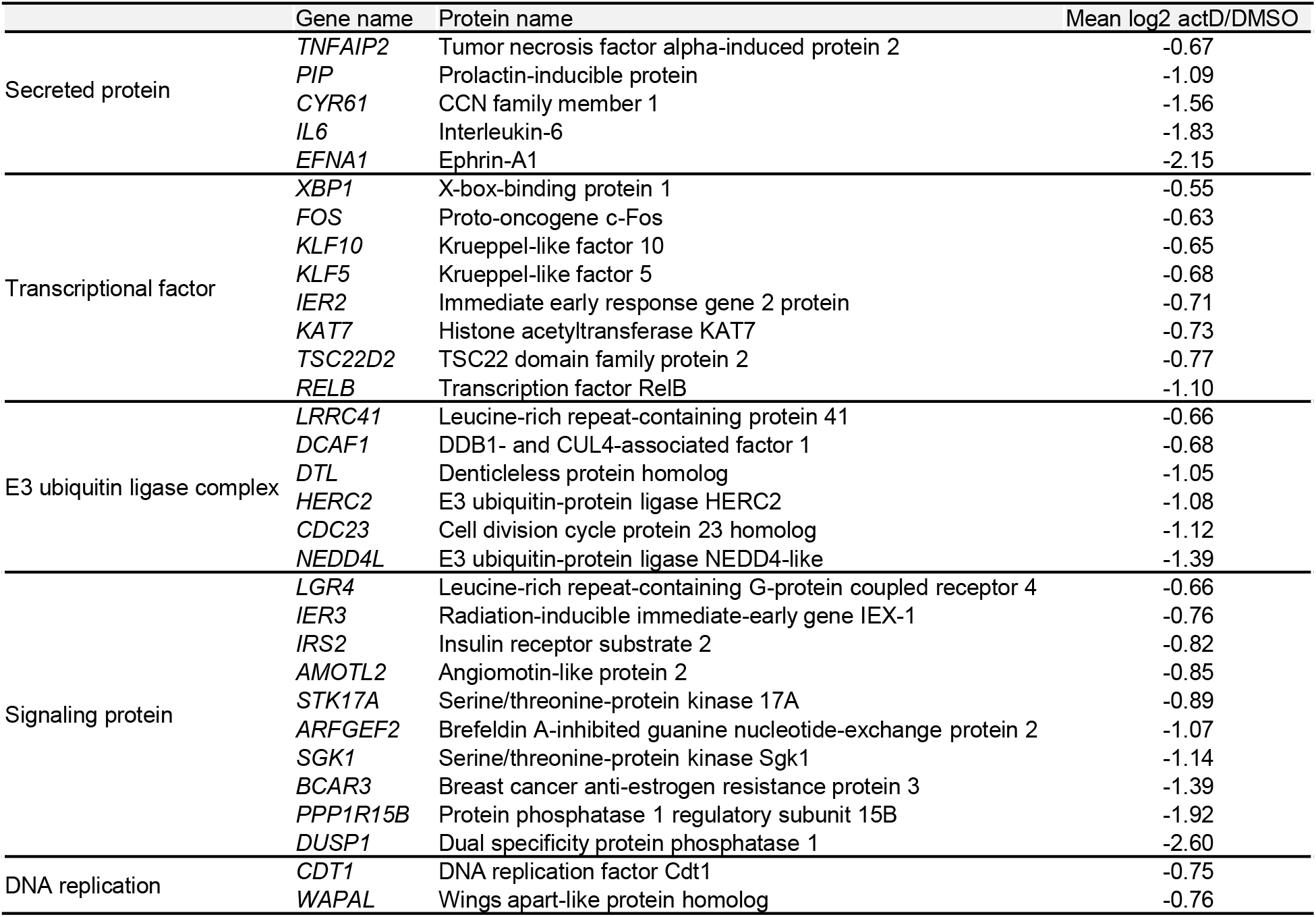
A list of selected proteins exclusively identified by the present method (related to Fig. 3C and Fig. 4B).

## Discussion

Our method presented here provides accurate quantitative profiles for thousands of NPCs that represent acute translation products, a key advantage over conventional methods (2–5) in which relatively longitudinal proteostasis and already-synthesized proteins (but not NPCs) can mainly be captured. Despite its advantage, it is also important to note the limitations of this method. First, affinity purification commonly involves non-specific protein binding to beads, which hampers an accurate and precise identification and quantification of NPCs. In this context, we observed that lower intensities of NPCs in comparison to non-specific proteins could compromise reproducibility (Supplementary Fig. 1B, 1C and 2B). Second, the OPP treatment may induce a secondary effect during the labeling (28). Third, the experimental procedure involves multiple steps, including pulse labeling, click reaction, and affinity purification, in addition to a conventional proteomic workflow. Fourth, the use of OPP and SILAC amino acids is more expensive than normal puromycin and amino acids.

Our data showed that RPs were among the most regulated proteins upon actD treatment (Fig. 3C). Although it is not surprising that the inhibition of rRNA synthesis suppressed the transcription/translation of RP-encoding genes, we showed that only a small subset of the RPs such as RPLP0, RPLP1, RPL12, RPL13, and RPL24 was modulated (Fig. 3C). Why are RPs differentially regulated? RPLP0, RPLP1, and RPL12 are known to form the ribosomal P-stalk which is built at the final step of the ribosome maturation in the cytoplasm (29). A previous pulse-labeling experiment also showed that newly synthesized RPLP0, RPL12, RPL13, and RPL24 are subjected to slow or not exported to the nucleus (30). These studies collectively indicate that the specific RPs may be loaded on the pre-ribosomal complex, possibly in the cytoplasm, during the last steps of ribosome maturation. It is therefore tempting to speculate that for maintaining ribosome biogenesis at the multiple levels, the abundance level of the RP subset is regulated at the level of protein synthesis in the cytoplasm, whereas the proteostasis of other RPs is maintained in nucleoli. In line with this notion, the other regulated RPs, RPS9, RPS23, and RPLP1, were also implicated as those assembled during the late stage of ribosome complex formation (Supplementary Fig. 2C) (31).

While actD does not affect translation itself (32), our results in Fig. 3C may reflect direct and/or indirect consequences upon the inhibition of RNAPs. For example, it is known that transcription inhibitors, including actD, cause enhanced transcription of a specific set of genes due to feedback loops (33, 34), p53 accumulation leading to apoptosis (35), and changes in the nuclear structure (36). Further investigation is required, particularly by combining RNA sequencing to quantify the mRNA level, to dissect the underlying transcriptional and post-transcriptional regulation.

In conclusion, we demonstrated that the present approach is a powerful tool to profile acute translational regulation at the NPC level. Thus, it can be applied to many biological contexts such as stress responses and virus infection. Further improvements towards higher deep proteome coverage, a simpler protocol and feasibility for *in vivo* systems will make this method more advantageous.

## Abbreviations

LC/MS/MS: liquid chromatography-tandem mass spectrometry
OPP: O-propargyl-puromycin
SILAC: stable isotope labeling of amino acids in cell culture
QuanCAT: quantitative noncanonical amino acid tagging
NPC: nascent polypeptide chain
PUNCH-P: puromycin-associated nascent chain proteomics
CHX: cycloheximide
iBAQ: intensity-based absolute quantification
actD: actinomycin D
RP: ribosomal protein
RPF: ribosome protected fragment
RNAP: RNA polymerase

## Acknowledgements

We thank the members of the Department of Molecular & Cellular BioAnalysis for fruitful discussion. KI thanks the Samuro Kakiuchi Memorial Research Award for Young Scientists for supporting this study.

---

**Supplementary Figure 1.**
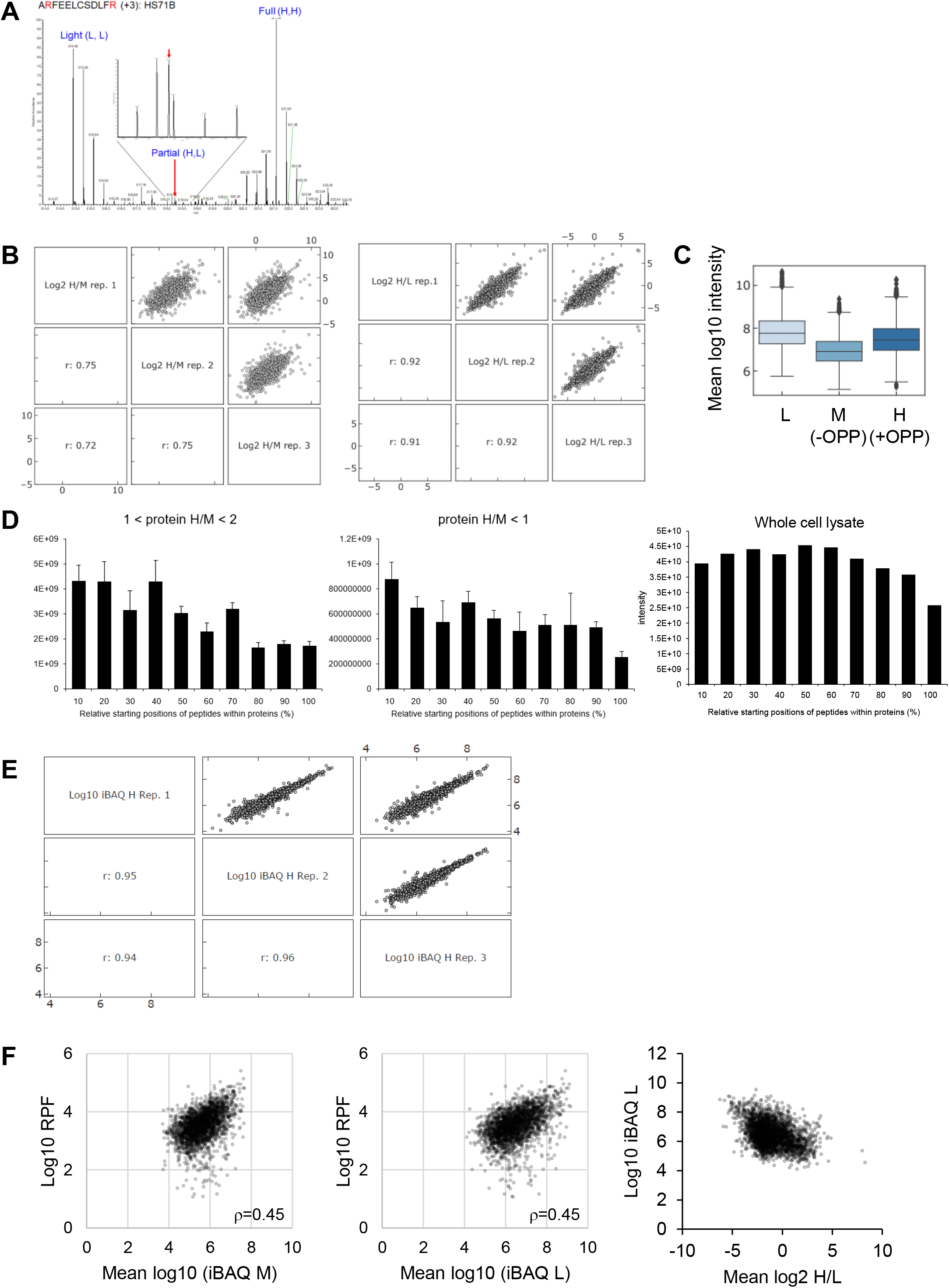
Systematic evaluation of the method. (A) The effect of “light” amino acids on pulse SILAC labeling. (B) Multi scatter plots showing log2 H/M (left) and log2 H/L (right). Pearson correlation coefficient (r) was shown. (C) A box plot showing mean log10 intensities of non-specific proteins (“L”), OPP(-) control (“M”) and OPP(+) (“H”). (D) Relative starting positions of identified peptides within proteins. Proteins that exhibit 1<H/M<2 (left) and H/M<1 (center) were shown. As a control, peptides identified from a HeLa whole lysate was also shown (right). (E) A multi scatter plot showing log10 iBAQ “H” intensity for only proteins that exhibited H/M>2. (F) Scatter plots showing the correlation between average iBAQ (“M”: left, “L”: right) and ribosome footprint read counts. A scatter plot showing the correlation between average iBAQ “L” and average log2 H/L (right). These results indicate that the degree to which proteins bind to beads non-specifically depends on their cellular protein abundance.

**Supplementary Figure 2.**
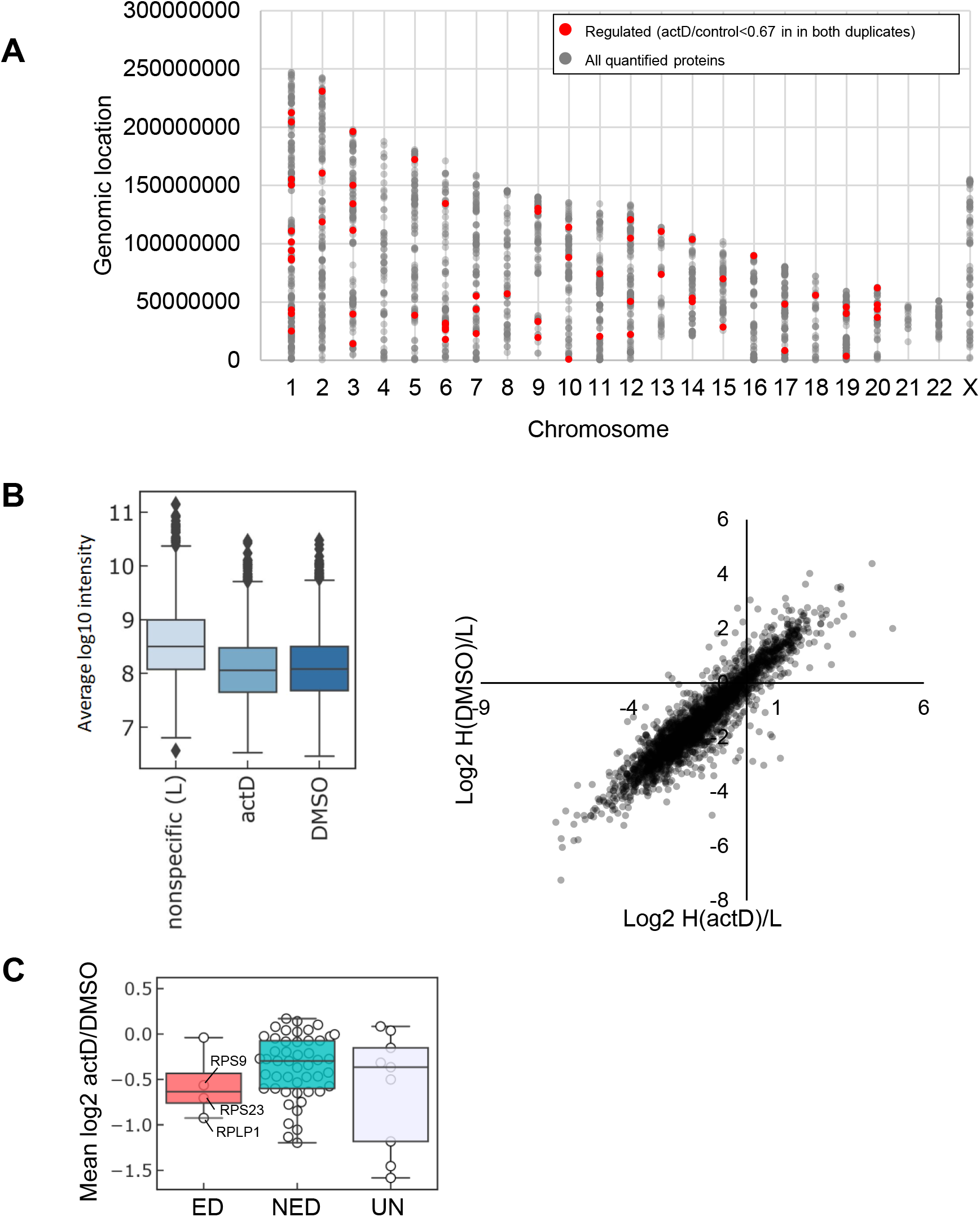
Systematic evaluation of nascent proteome in response to actinomycin D. (A) Genomic regions for proteins regulated by actD treatment (red) and all quantified proteins in this study (grey). (B) A box plot showing mean log10 intensities of non-specific proteins, actD- and DMSO-treated proteins (left). A scatter plot shows a higher correlation between log2 H/L than log2 H/M (right, see also Fig. 3C showing correlation between H/M). These results indicate that poor reproducibility of H/M ratios can be explained by the lower signal intensities of “H” and “M” proteins compared with “L” proteins. (C) Characterization of RPs. RPs were grouped into three categories; 1) exponentially degradation (ED) proteins, 2) non-exponentially degradation (NED) proteins and 3) undefined proteins, based on previous analysis on the proteome-wide degradation kinetics (30). It was shown that the ED proteins are prone to be assembled later than the NED proteins, and thus the ED ribosomal proteins may be assembled later than NED.

**Supplementary Table 1** (related to Fig. 2C): A list of proteins quantified (H: +OPP, M: -OPP).

**Supplementary Table 2** (related to Fig. 3C and Fig. 4B): A list of proteins quantified (actD vs DMSO).

## References

1. Truitt, M.L. and Ruggero, D. (2017) New frontiers in translational control of the cancer genome. Nat. Rev. Cancer, 17, 332.

2. Selbach, M., Schwanhäusser, B., Thierfelder, N., Fang, Z., Khanin, R. and Rajewsky, N. (2008) Widespread changes in protein synthesis induced by microRNAs. Nature, 455, 58–63.

3. Imami, K., Milek, M., Bogdanow, B., Yasuda, T., Kastelic, N., Zauber, H., Ishihama, Y., Landthaler, M. and Selbach, M. (2018) Phosphorylation of the Ribosomal Protein RPL12/uL11 Affects Translation during Mitosis. Mol. Cell, 72, 84–98.e9.

4. Howden, A.J.M., Geoghegan, V., Katsch, K., Efstathiou, G., Bhushan, B., Boutureira, O., Thomas, B., Trudgian, D.C., Kessler, B.M., Dieterich, D.C., et al. (2013) QuaNCAT: quantitating proteome dynamics in primary cells. Nat. Methods, 10, 343–346.

5. Eichelbaum, K., Winter, M., Berriel Diaz, M., Herzig, S. and Krijgsveld, J. (2012) Selective *enrichment of newly synthesized proteins for quantitative secretome analysis*. Nat. Biotechnol., 30, 984–990.

6. Dieterich, D.C., Link, A.J., Graumann, J., Tirrell, D.A. and Schuman, E.M. (2006) Selective identification of newly synthesized proteins in mammalian cells using bioorthogonal noncanonical amino acid tagging (BONCAT). Proc. Natl. Acad. Sci. U. S. A., 103, 9482–9487.

7. Pestka, S. (1971) Inhibitors of ribosome functions. Annu. Rev. Microbiol., 25, 487–562.

8. Aviner, R., Geiger, T. and Elroy-Stein, O. (2013) Novel proteomic approach (PUNCH-P) reveals cell cycle-specific fluctuations in mRNA translation. Genes Dev., 27, 1834–1844.

9. Forester, C.M., Zhao, Q., Phillips, N.J., Urisman, A., Chalkley, R.J., Oses-Prieto, J.A., Zhang, L., Ruggero, D. and Burlingame, A.L. (2018) Revealing nascent proteomics in signaling pathways and cell differentiation. Proc. Natl. Acad. Sci. U. S. A., 115, 2353–2358.

10. Barrett, R.M., Liu, H.-W., Jin, H., Goodman, R.H. and Cohen, M.S. (2016) Cell-specific Profiling *of Nascent Proteomes Using Orthogonal Enzyme-mediated Puromycin Incorporation*. ACS Chem. Biol., 11, 1532–1536.

11. Mellacheruvu, D., Wright, Z., Couzens, A.L., Lambert, J.-P., St-Denis, N.A., Li, T., Miteva, Y.V., Hauri, S., Sardiu, M.E., Low, T.Y., et al. (2013) The CRAPome: a contaminant repository for affinity purification-mass spectrometry data. Nat. Methods, 10, 730–736.

12. Imami, K., Sugiyama, N., Tomita, M. and Ishihama, Y. (2010) Quantitative proteome and phosphoproteome analyses of cultured cells based on SILAC labeling without requirement of serum dialysis. Mol. Biosyst., 6, 594–602.

13. Adachi, J., Hashiguchi, K., Nagano, M., Sato, M., Sato, A., Fukamizu, K., Ishihama, Y. and Tomonaga, T. (2016) Improved Proteome and Phosphoproteome Analysis on a Cation Exchanger by a Combined Acid and Salt Gradient. Anal. Chem., 88, 7899–7903.

14. Ishihama, Y., Rappsilber, J., Andersen, J.S. and Mann, M. (2002) Microcolumns with self-assembled particle frits for proteomics. J. Chromatogr. A, 979, 233–239.

15. Cox, J. and Mann, M. (2008) MaxQuant enables high peptide identification rates, *individualized p.p.b.-range mass accuracies and proteome-wide protein quantification*. Nat. Biotechnol., 26, 1367–1372.

16. Cox, J., Neuhauser, N., Michalski, A., Scheltema, R.A., Olsen, J.V. and Mann, M. (2011) Andromeda: a peptide search engine integrated into the MaxQuant environment. J. Proteome Res., 10, 1794–1805.

17. Nolte, H., MacVicar, T.D., Tellkamp, F. and Krüger, M. (2018) Instant Clue: A Software Suite for Interactive Data Visualization and Analysis. Sci. Rep., 8, 12648.

18. Okuda, S., Watanabe, Y., Moriya, Y., Kawano, S., Yamamoto, T., Matsumoto, M., Takami, T., Kobayashi, D., Araki, N., Yoshizawa, A.C., et al. (2017) jPOSTrepo: an international standard data repository for proteomes. Nucleic Acids Res., 45, D1107–D1111.

19. Moriya, Y., Kawano, S., Okuda, S., Watanabe, Y., Matsumoto, M., Takami, T., Kobayashi, D., Yamanouchi, Y., Araki, N., Yoshizawa, A.C., et al. (2019) The jPOST environment: an integrated proteomics data repository and database. Nucleic Acids Res., 47, D1218–D1224.

20. Schwanhäusser, B., Busse, D., Li, N., Dittmar, G., Schuchhardt, J., Wolf, J., Chen, W. and Selbach, M. (2011) Global quantification of mammalian gene expression control. Nature, 473, 337–342.

21. Stumpf, C.R., Moreno, M.V., Olshen, A.B., Taylor, B.S. and Ruggero, D. (2013) The Translational Landscape of the Mammalian Cell Cycle. Mol. Cell, 52, 574–582.

22. Shuda, M., Velásquez, C., Cheng, E., Cordek, D.G., Kwun, H.J., Chang, Y. and Moore, P.S. (2015) CDK1 substitutes for mTOR kinase to activate mitotic cap-dependent protein translation. Proc. Natl. Acad. Sci. U. S. A., 112, 5875–5882.

23. Imami, K. and Yasuda, T. (2019) Measuring Protein Synthesis during Cell Cycle by Azidohomoalanine (AHA) Labeling and Flow Cytometric Analysis. BIO-PROTOCOL, 9.

24. Ingolia, N.T., Ghaemmaghami, S., Newman, J.R.S. and Weissman, J.S. (2009) Genome-wide analysis in vivo of translation with nucleotide resolution using ribosome profiling. Science, 324, 218–223.

25. Kohler, R., Mooney, R.A., Mills, D.J., Landick, R. and Cramer, P. (2017) Architecture of a transcribing-translating expressome. Science, 356, 194–197.

26. Perry, R.P. and Kelley, D.E. (1970) Inhibition of RNA synthesis by actinomycin D: Characteristic dose-response of different RNA species. Journal of Cellular Physiology, 76, 127–139.

27. Bensaude, O. (2011) Inhibiting eukaryotic transcription: Which compound to choose? How to evaluate its activity? Transcription, 2, 103–108.

28. Marciano, R., Leprivier, G. and Rotblat, B. (2018) Puromycin labeling does not allow protein synthesis to be measured in energy-starved cells. Cell Death Dis., 9, 39.

29. Lo, K.-Y., Li, Z., Wang, F., Marcotte, E.M. and Johnson, A.W. (2009) Ribosome stalk assembly *requires the dual-specificity phosphatase Yvh1 for the exchange of Mrt4 with P0*. J. Cell Biol., 186, 849–862.

30. Lam, Y.W., Lamond, A.I., Mann, M. and Andersen, J.S. (2007) Analysis of nucleolar protein dynamics reveals the nuclear degradation of ribosomal proteins. Curr. Biol., 17, 749–760.

31. McShane, E., Sin, C., Zauber, H., Wells, J.N., Donnelly, N., Wang, X., Hou, J., Chen, W., Storchova, Z., Marsh, J.A., et al. (2016) Kinetic Analysis of Protein Stability Reveals Age-Dependent Degradation. Cell, 167, 803–815.e21.

32. Cooper, H.L. and Braverman, R. (1977) The mechanism by which actinomycin D inhibits protein synthesis in animal cells. Nature, 269, 527–529.

33. Cassé, C., Giannoni, F., Nguyen, V.T., Dubois, M.F. and Bensaude, O. (1999) The transcriptional inhibitors, actinomycin D and alpha-amanitin, activate the HIV-1 promoter *and favor phosphorylation of the RNA polymerase II C-terminal domain*. J. Biol. Chem., 274, 16097–16106.

34. Imamichi, T., Conrads, T.P., Zhou, M., Liu, Y., Adelsberger, J.W., Veenstra, T.D. and Lane, H.C. (2005) A transcription inhibitor, actinomycin D, enhances HIV-1 replication through an interleukin-6-dependent pathway. J. Acquir. Immune Defic. Syndr., 40, 388–397.

35. Galluzzi, L., Morselli, E., Kepp, O., Vitale, I., Pinti, M. and Kroemer, G. (2011) Mitochondrial liaisons of p53. Antioxid. Redox Signal., 15, 1691–1714.

36. Boulon, S., Westman, B.J., Hutten, S., Boisvert, F.-M. and Lamond, A.I. (2010) The nucleolus under stress. Mol. Cell, 40, 216–227.

